# Software tools for visualizing Hi-C data

**DOI:** 10.1101/086017

**Authors:** Galip Gürkan Yardımcı, William Stafford Noble

## Abstract

Recently developed, high-throughput assays for measuring the three-dimensional configuration of DNA in the nucleus have provided unprecedented insights into the relationship between DNA 3D configuration and function. However, accurate interpretation of data from assays such as ChIA-PET and Hi-C is challenging because the data is large and cannot be easily rendered using a standard genome browser. In particular, an effective Hi-C visualization tool must provide a variety of visualization modes and be capable of viewing the data in conjunction with existing, complementary data. We review a number of such software tools that have been described recently in the literature, focusing on tools that do not require programming expertise on the part of the user. In particular, we describe HiBrowse, Juicebox, my5C, the 3D Genome Browser, and the Epigenome Browser, outlining their complementary functionalities and highlighting which types of visualization tasks each tool is best designed to handle.

## Introduction

The three-dimensional conformation of the genome in the nucleus influences multiple key biological processes, including transcriptional regulation and DNA replication timing. Over the past decade, a series of chromosome conformation capture assays have been developed for characterizing 3D contacts associated with a single locus (3C, 4C) [1–3], a set of loci (5C, ChIA-PET) [4, 5] or the whole genome (Hi-C) [6]. Using these assays, researchers have profiled the conformation of chromatin in a variety of organisms and systems, revealing a hierarchical, domain-like organization of chromatin.

Here we focus on the Hi-C assay and variants thereof, which provide a genome-wide view of chromosome conformation. The assay consists of five steps: (1) crosslinking of DNA with formaldehyde, (2) cleaving cross-linked DNA with an endonuclease, (3) ligating the ends of cross-linked fragments to form a circular molecule marked with biotin, (4) shearing circular DNA and pulling down fragments marked with biotin, and (5) paired-end sequencing of the pulled-down fragments. A pair of sequence reads from a single ligated molecule map to two distinct regions of genome, and the abundance of such fragments provides a measure of how frequently, within a population of cells, the two loci are in contact. Thus, in contrast to assays such as DNase-seq and ChIP-seq [7, 8], which yield a one-dimensional count vector across the genome, the output of Hi-C is a two-dimensional matrix of counts, with one entry for each pair of genomic loci. In practice, production of this matrix involves a series of filtering and normalization steps (reviewed in [9] and [10]).

A critical parameter in any Hi-C analysis pipeline is the effective resolution at which the data is analyzed [10, 11]. In this context, “resolution” simply refers to the size of the loci for which Hi-C counts are aggregated. Using current technology, sequencing deeply enough to achieve very high resolution data for large genomes is prohibitively expensive. In principle, a basepair resolution analysis of the human genome would require aggregating counts across a matrix of size approximately (3 × 10^9^)^2^ = 9 × 10^18^. In practice, reads that fall within a contiguous genomic window are binned together, reducing the size and sparsity of the matrix at the cost of resolution. Following this process, Hi-C data can be represented as a “contact matrix” *M*, where entry *M*_*ij*_ is the number of Hi-C read pairs, or contacts, between genomic locations designated by bin *i* and bin *j*.

For researchers studying chromatin conformation, Hi-C data presents a number of significant analytical challenges. Filtering and normalization strategies can be employed to correct experimental artifacts and biases [9–11]. Statistical confidence measures can be estimated to identify sets of high confidence contacts [12]. The observed Hi-C data can be compared with and correlated against complementary data sets measuring protein-DNA interactions, gene expression, and replication timing [13–15]. And the 3D conformation of the DNA itself can be estimated from the Hi-C data, potentially taking into account data derived from other assays or from multiple experimental conditions [16–19]

Complementary to these analyses is the ability to efficiently and accurately visualize Hi-C data. Such visualizations are quite challenging, both because Hi-C data is large and because many existing tools for visualizing large-scale genomic data, such as genome browsers, do not directly generalize to visualizing data defined over pairs of loci [20, 21]. Furthermore, many biological hypotheses involve multiple biological processes and hence require jointly visualizing Hi-C data with other chromatin features. Thus, visualizing Hi-C data alone is not enough—a truly useful tool must integrate a variety of different types of genomic data and annotations.

To address these challenges, a variety of software tools have been described recently that provide robust and informative methods for making sense out of Hi-C data. Here we investigate five tools that can be operated using a web browser or a graphical user interface: Hi-Browse [22], my5C [23], Juicebox [24], the Epigenome Browser [25] and the 3D Genome Browser (http://www.3dgenome.org) (Table 1). These tools require no programming skills and are thus useful to a wide audience of researchers. We assess these tools using multiple criteria, such as the types of visualizations provided by the tool, the ability to integrate multiple visualization modes, and the number and variety of available datasets in a given tool. In particular, we note the suitability of each tool to different types of inquiry regarding the 3D structure of the genome and its interplay with other biological processes. We present examples that range from large scale visualizations of Hi-C data from whole genomes and chromosomes to fine scale local visualizations of putative promoter enhancer interactions and DNA loops, highlighting additional tools-specific capabilities that complement each visualization type.

**Table 1:**
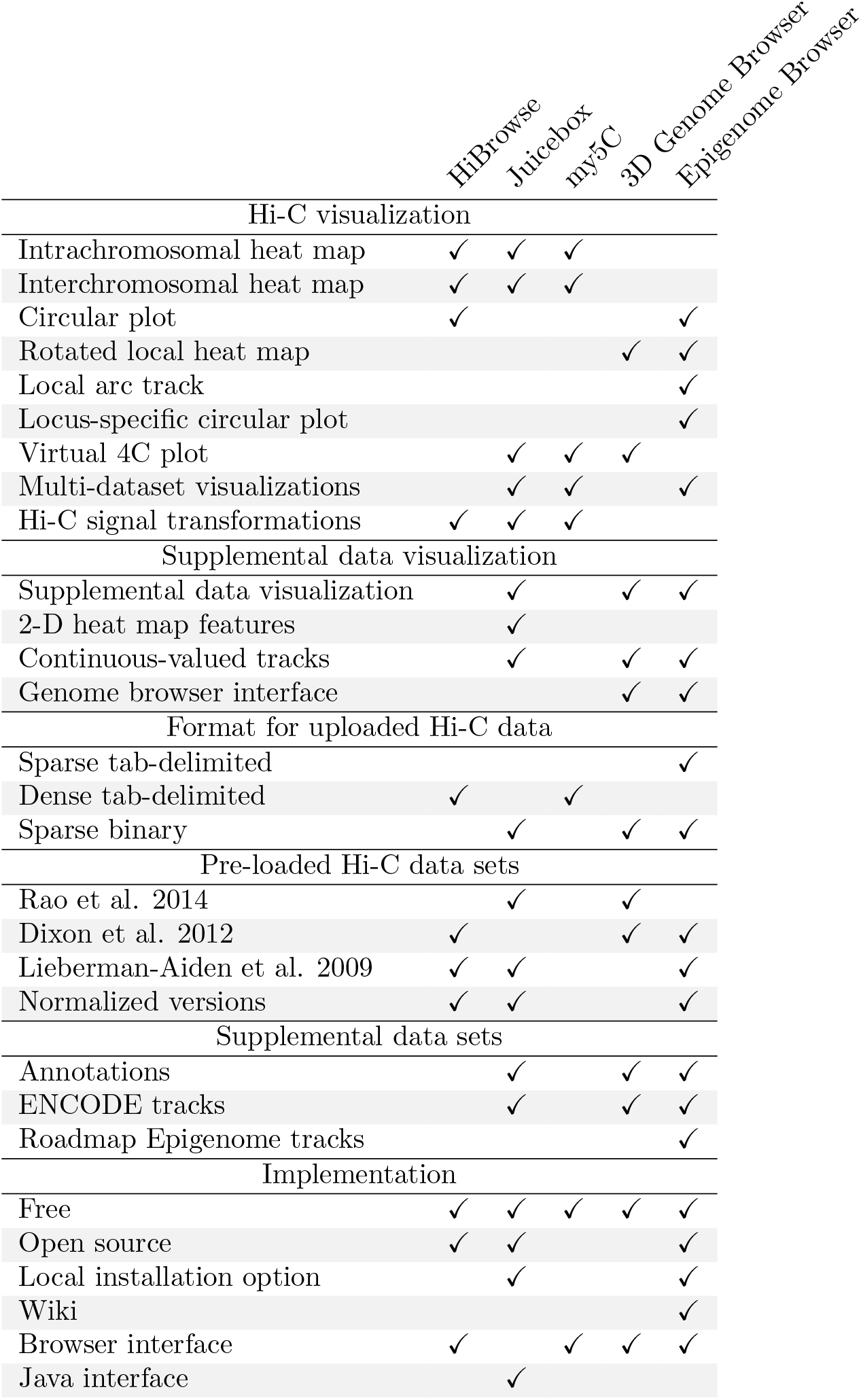
Comparison of toolkit functionality.

|  | HiBrowse | Juicebox | my5C | 3D Genome Browser | Epigenome Browser |
| --- | --- | --- | --- | --- | --- |
| Hi-C visualization |  |  |  |  |  |
| Intrachromosomal heat map | ✓ | ✓ | ✓ |  |  |
| Interchromosomal heat map | ✓ | ✓ | ✓ |  |  |
| Circular plot | ✓ |  |  |  | ✓ |
| Rotated local heat map |  |  |  | ✓ | ✓ |
| Local arc track |  |  |  |  | ✓ |
| Locus-specific circular plot |  |  |  |  | ✓ |
| Virtual 4C plot |  | ✓ | ✓ | ✓ |  |
| Multi-dataset visualizations |  | ✓ | ✓ |  | ✓ |
| Hi-C signal transformations | ✓ | ✓ | ✓ |  |  |
| Supplemental data visualization |  |  |  |  |  |
| Supplemental data visualization |  | ✓ |  | ✓ | ✓ |
| 2-D heat map features |  | ✓ |  |  |  |
| Continuous-valued tracks |  | ✓ |  | ✓ | ✓ |
| Genome browser interface |  |  |  | ✓ | ✓ |
| Format for uploaded Hi-C data |  |  |  |  |  |
| Sparse tab-delimited |  |  |  |  | ✓ |
| Dense tab-delimited | ✓ |  | ✓ |  |  |
| Sparse binary |  | ✓ |  | ✓ | ✓ |
| Pre-loaded Hi-C data sets |  |  |  |  |  |
| Rao et al. 2014 |  | ✓ |  | ✓ |  |
| Dixon et al. 2012 | ✓ |  |  | ✓ | ✓ |
| Lieberman-Aiden et al. 2009 | ✓ | ✓ |  |  | ✓ |
| Normalized versions | ✓ | ✓ |  |  | ✓ |
| Supplemental data sets |  |  |  |  |  |
| Annotations |  | ✓ |  | ✓ | ✓ |
| ENCODE tracks |  | ✓ |  | ✓ | ✓ |
| Roadmap Epigenome tracks |  |  |  |  | ✓ |
| Implementation |  |  |  |  |  |
| Free | ✓ | ✓ | ✓ | ✓ | ✓ |
| Open source | ✓ | ✓ |  |  | ✓ |
| Local installation option |  | ✓ |  |  | ✓ |
| Wiki |  |  |  |  | ✓ |
| Browser interface | ✓ |  | ✓ | ✓ | ✓ |
| Java interface |  | ✓ |  |  |  |

## Large scale visualization

The three-dimensional conformation of a complete chromosome or genome is most commonly visualized by one of two different methods. The contact matrix can be represented as a square heat map, where the color corresponds to the contact count, or the genome can be represented as a circle, with contacts indicated by edges connecting distal pairs of loci. In principle, other types of large-scale visualizations are feasible, using for example a graph with nodes as loci and edges as contacts, but these have not proved as useful as heat maps and circular plots.

Between these two visualization schemes, perhaps the most straightforward for a Hi-C contact matrix is a heat map. Contact matrices are by definition symmetric around the diagonal, and the number of rows and columns is equal to the length of the genome divided by the bin size. The color scale associated with the heat map may correspond to raw contact counts or counts that have been appropriately normalized. The dominant visual feature in every Hi-C heat map is the strong diagonal, which represents the 3D proximity of pairs of loci that are adjacent in genomic coordinates. Heat maps can be constructed for the full genome (Figure 1A) or for individual chromosomes (Figure 1B). Low resolution (1–10 MB resolution) contact matrices are typically sufficient for full genome visualizations and can be produced, for the human genome, using Hi-C datasets containing tens of millions of read pairs. Whole genome visualizations can reveal potential rearragements of the genome (Figure 1A), whereas single chromosome visualizations are useful in identifying large-scale properties of chromatin conformation, such as chromosome compartments or the bipartite structure of the mouse inactive X chromosome (Figure 1B). Three of the five tools that we investigated—HiBrowse, Juicebox, and my5C—provide heat map visualizations.

**Figure 1:**
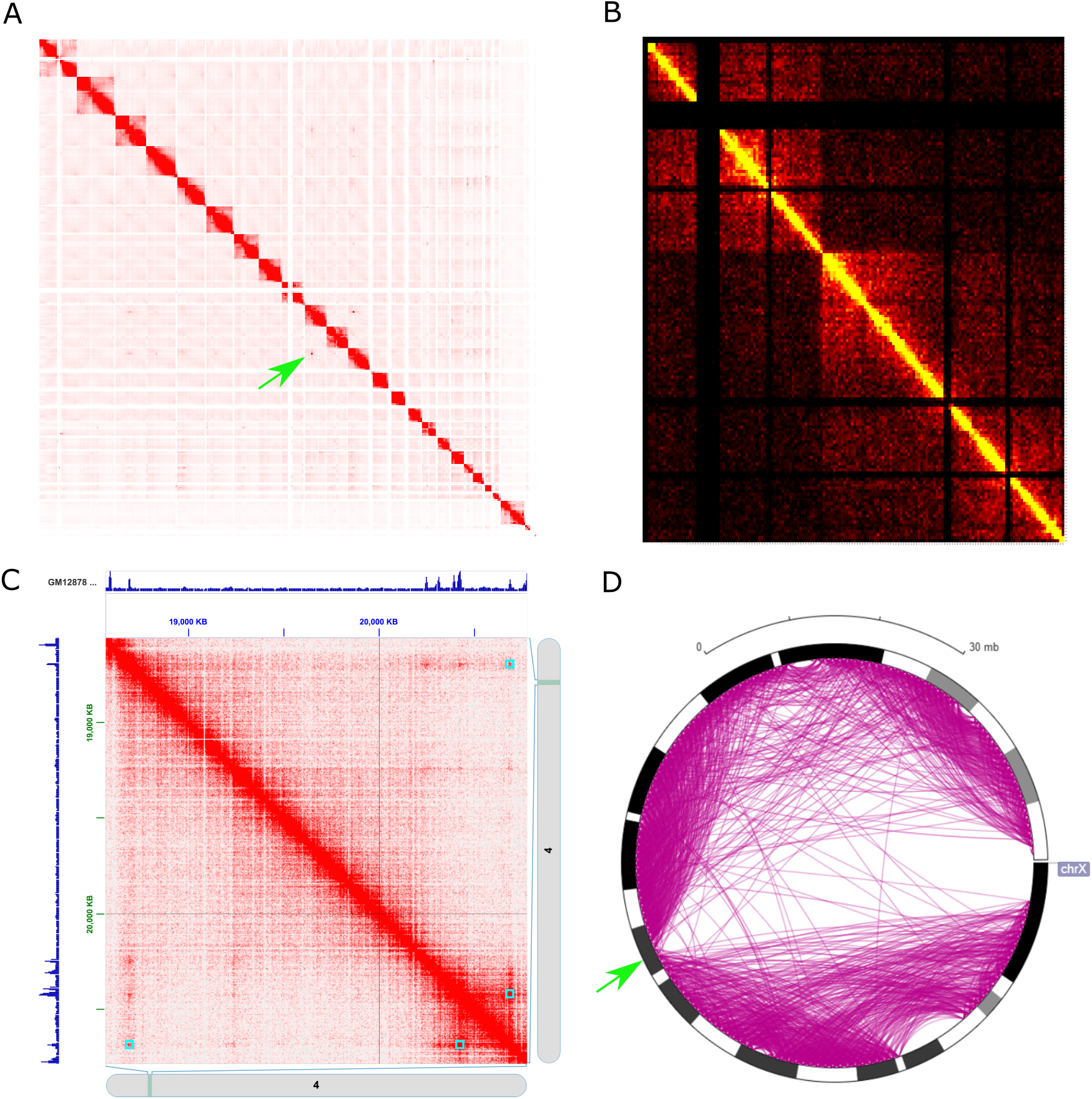
Heat map and circular plot visualization of Hi-C data. (A) Hi-C interactions among all chromosomes from G401 human kidney cells, as plotted by my5C. The green arrow points to aberrant interchromosomal signal in the Hi-C matrix, possibly caused by a rearragement event. (B) Heat map visualization illustrating the bipartite structure of the mouse X chromosome, as plotted by Hi-Browse, using in-situ DNase Hi-C data [26]. (C) Heat map visualization of a 3 Mbp locus (chr4:18000000-21000000) reveals the presence of loops that coincide with CTCF binding sites, validated by CTCF peaks shown on the top and left of the heat map. Computationally annotated loops are displayed as blue squares in the heat map. This heat map was produced by Juicebox, using in-situ Hi-C data from the GM12878 cell line [27]. (D) Circular plot of the bipartite mouse X chromosome, showing a striking depletion of arcs between the two mega-domains, the locus separating the mega-domains is shown by a green arrow. The plot was generated by the Epigenome Browser.

A heat map can also be used to visualize the conformation of a locus of interest. This setting zooms into a region of the full contact matrix, visualized at higher resolution. The resulting map can be useful to visually identify loops, i.e., distal regions of DNA that exhibit unusually high contact counts relative to neighboring pairs of loci. Loop annotations detected by loop-finding algorithms can be displayed directly on a Hi-C contact map by Juicebox. Loop formation has been shown to depend on DNA binding of the CTCF protein [28]; therefore, it is desirable to jointly visualize CTCF binding data from a ChIP-seq assay alongside Hi-C data, to assist in the interpretation of possible loops. Juicebox can plot data from other assays or genomic features, either as binary features or continuous signal plots, placing them on the sides of heat map (Figure 1C).

Circular plots, originally designed to visualize genomic data, offer an alternative way to visualize Hi-C data at chromosome scale. In this type of plot, the circle typically represents the full length of a chromosome, and Hi-C contacts are represented by arcs (Figure 1D). It is straightforward to convert a contact matrix into a circular plot: loci *i* and *j* are connected by an arc if entry *M*_*ij*_ in the contact matrix exceeds a user-specified cutoff value. Hi-Browse and the Epigenome Browser can both generate circular plots.

## Local visualization

Although Hi-C data spans the full genome, many hypotheses require closely inspecting a particular region or regions of interest. A common way to visualize multiple genomic data sets at a particular locus is via a genome browser, in which the DNA is arrayed horizontally and various types of data appear in parallel with the DNA sequence. Two of the tools that we investigated, the 3D Genome Browser and the Epigenome Browser, extend the browser framework to incorporate Hi-C data, thereby offering rich and complex representations of DNA sequence, chromatin, gene structure, regulatory elements and 3D conformation.

Four different visualization modes are available in the context of a genome browser. Both the 3D Genome Browser and the Epigenome Browser provide a heat map visualization, in which the upper triangle of the contact matrix is rotated by 45 degrees and then aligned so that the bins of the matrix correspond to chromosomal coordinates (Figure 2A). This heat map visualization is limited to capturing intra-chromosomal contacts, and the genomic distance between contacts is limited by the vertical screen space available to the heat map track. This makes displaying distal contacts at high resolution impractical.

**Figure 2:**
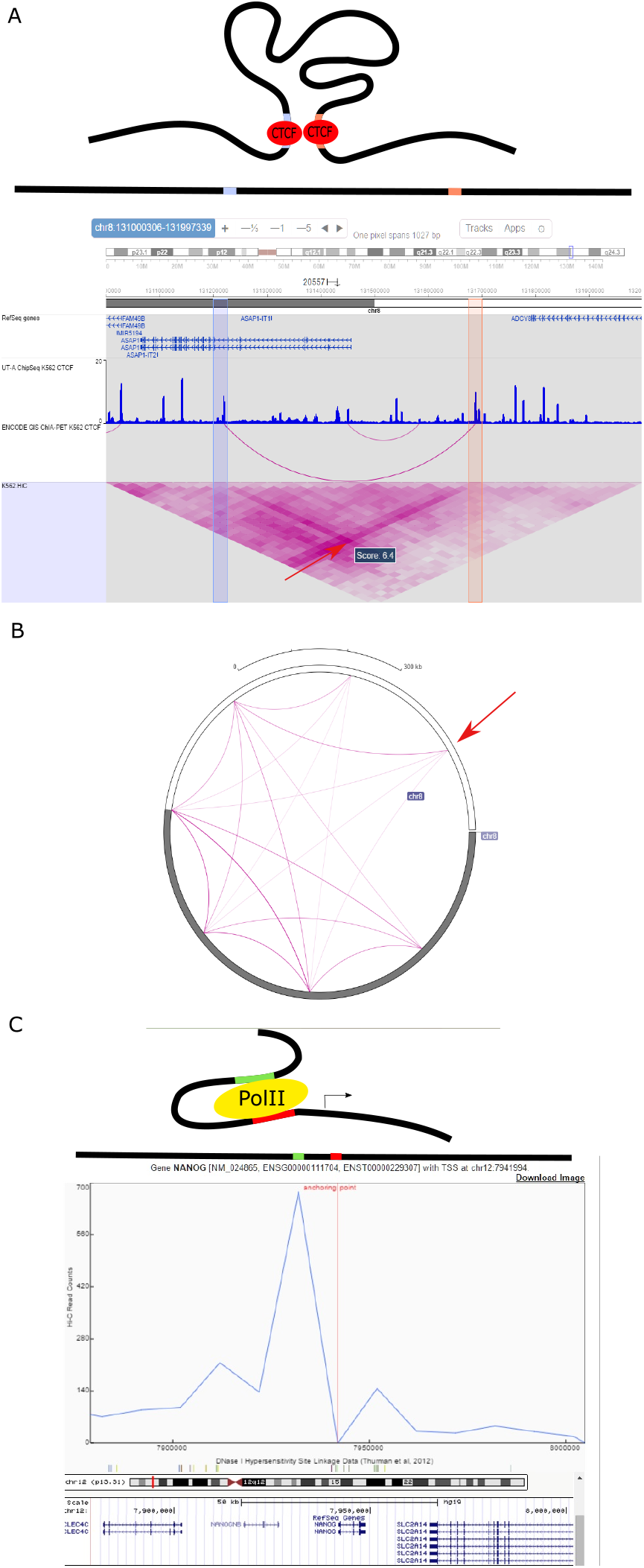
Local visualization modes. (A) A cartoon representation of the 3D conformation of a putative DNA loop tethered by two CTCF proteins. CTCF binding sites are colored in blue and pink on the black DNA strand. Below the cartoon, a 1D representation of the DNA fragment that forms the loop is placed above an Epigenome Browser visualization of a ~1MB locus, displaying the genes, CTCF binding, and interactions detected by ChIP-seq [29] and ChIA-PET assays (unpublished data, GEO ID:GSM970216), as well as 3D interactions as measured by Hi-C [27]. Two bins containing putative binding sites (pink and blue bars) show an enrichment of Hi-C contacts in the heatmap visualization [27] (indicated by the red arrow). CTCF tethered interactions measured by ChIA-PET in an arcs view also indicate an interaction between these two putative binding sites. (B) A circular plot displaying the chromosome-wide long range contacts of the CTCF loop in panel A; the locus of interest is highlighted by a red arrow. The contacts are displayed as arcs, and only contacts above a certain threshold are visualized. (C) A putative promoter-enhancer interaction around the NANOG gene is displayed as a cartoon, including the PolII complex (yellow oval). Red and green bars in these cartoons represent the promoter and enhancer elements, respectively. Below the cartoon representations, a virtual 4C plot from the 3D genome browser is shown, visualizing the Hi-C signal around NANOG promoter with a 1D representation of this region aligned above the plot. The bin in focus (the “anchoring point”) corresponds to the promoter of the NANOG gene. The height of the blue line indicates, for each locus, the read count for contacts between the current locus and the anchor point. In particular, the series shows an upstream enrichment of signal from a capture Hi-C experiment specifically targeting the NANOG promoter [30], suggesting a promoter-enhancer interaction. This observation is further supported by enrichment of DNaseI linkage data [31] (shown in grey below the primary plot) around the promoter and upstream regions. The NANOG gene is shown in the UCSC Genome Browser track under the virtual 4C plot.

A second type of Hi-C visualization is a local arc track, also available in both the 3D Genome and Epigenome Browsers (Figure 2A). Similar to a circular plot, a local arc track connects two genomic loci with an arc if the corresponding Hi-C signal is above a user specified threshold. Compared to heat map tracks, arc tracks offer a simpler interpretation of Hi-C contacts, at the expense of leaving out some of the data. The Epigenome Browser can display both Hi-C and ChIA-PET interactions in arc view, while the 3D Genome Browser uses arc tracks exclusively for ChIA-PET interactions.

The Epigenome Browser offers a third visualization mode that is intermediate between a local and global view. This is a global circular plot that includes contacts between a selected locus, pointed by a red arrow in (Figure 2B) and the rest of the genome or a single chromosome. This plots offers a simplified way of visualizing relevant long distance genome-wide contacts involving a specific locus.

Finally, the 3D Genome Browser offers a fourth visualization mode, called a virtual 4C plot, which is a slight modification of the local arc track (Figure 2C). Unlike a local arc track, which shows all contacts whose start and end loci are contained within the current browser view, a virtual 4C plot restricts the set of arcs to those involving a single user-specified locus. Thus, a virtual 4C plot for the locus corresponding to bin *i* is equivalent to plotting the entries from the *i*^*th*^ row of the contact matrix. By focusing on a single locus a virtual 4C plot is useful for testing specific hypotheses regarding the bin of interest. We also note that Juicebox and my5C offer a limited version of a 4C plot in the form a track alongside a heatmap visualization.

All of these local visualization modes are particularly useful within the context of a full genome browser where, for example, potential regulatory contacts can be easily inspected alongside gene annotations, histone ChIP-seq experiments that mark enhancers and promoters, etc. For example, the Epigenome Browser can provide a view of a potential CTCF-tethered loop alongside multiple tracks: gene annotations, Hi-C and ChIA-PET contacts and CTCF ChIP-seq signal (Figure 2A). The resulting visualization is a concise and rich representation of multiple types of data, increasing our confidence in the existence of this DNA loop.

## Data availability

Getting data into a Hi-C visualization tool can be accomplished in two ways: either the data is pre-loaded by the tool developers or the user is responsible for uploading their own data. Both modes of data entry can be provided in a single tool. Here we describe available data sets and upload capabilities for the five tools we investigated, including both Hi-C data sets and auxiliary genomic data sets.

## Hi-C datasets

The visualization tools we review all come with some publicly available datasets, with the exception of my5C. Available datasets include three influential studies that performed Hi-C experiments on multiple cell types, which we refer to via the last name of the first author on the respective publications: Lieberman-Aiden [6], Dixon [13], and Rao [27]. These three studies cover nine human cell types from different lineages and tissues—IMR90, H1, GM06990, HMEC, NHEK, K562, HUVEC, HeLa, and KBM7—making them useful for many types of analyses. In Table 1, we summarize the datasets available for each tool. In addition, Juicebox offers datasets from 27 additional studies, including data from a variety of organisms (Supplement Table 1). While most of these datasets are from Hi-C experiments performed on human cells, each tool supports genomes of other organisms. The Epigenome Browser supports a total of 19 genomes, and the 3D Genome browser supports human and mouse genomes. The other tools can be used with any genome.

Hi-C datasets are accumulating rapidly, and many users will need the capability to upload new datasets into these tools. All tools offer the ability to upload user data or data downloaded from repositories such as 3DGD [32] or 4DGenome [33]. Most tools accept files representing contact matrices; however, the file format requirements differ by tool (Table 1). The Epigenome Browser represents Hi-C matrices using tab-delimited text files, similar to the BED files commonly used in genomics. Hi-Browse and my5C also uses tab delimited text files, but unlike the Epigenome Browser format, the my5C and Hi-Browse formats require that every entry be explicitly represented in the input file, including pairs of loci with zero contacts. The 3D Genome Browser uses its own sparse matrix representation in binary format, which can be created using the BUTLRTools software package (https://github.com/yuelab/BUTLRTools). Juicebox depends on its accompanying tool Juicer [34] to build .hic files that store binary contact matrices at multiple resolutions. This .hic files are built from sequenced read pair files from a Hi-C experiment. The Epigenome Browser also supports the .hic format.

As Hi-C datasets continue to accumulate, the scientific community will likely come to a consensus on standardized file formats to represent Hi-C datasets. Most of the currently available file formats are very similar to one another, and conversion between most formats is straightforward using command line tools. An important tradeoff between different formats is the size of the file; sparse representations and especially the binary BUTLR and .hic formats require less disk space relative to uncompressed versions of other file formats.

## Data handling

Any given Hi-C data set can be binned at different resolutions. Generally, the user chooses a resolution value (i.e., bin size) based on the sequencing depth of the dataset, striking a balance between detail and the sparsity that results from high resolution analysis. All tools in this review support visualization of Hi-C matrices at various resolutions. Available datasets for each tool are stored at different resolution values, typically ranging from 1 Mb to as low as 5 kb. For user-uploaded datasets, it is up to the user to generate contact matrices at different resolutions, except for the .hic format which stores multiple resolutions in a single file.

After the resolution is set by the user, Hi-C data may be transformed in various ways to focus on different features of the data. The three most common transformations are matrix balancing to remove bin-specific biases [35–38], calculation of a correlation matrix for visualization of A and B compartments [6, 39], and calculating the ratio of observed over expected Hi-C counts to account for the so-called “genomic distance effect” (the density of interactions close to the diagonal in the Hi-C matrix) [6]. Hi-Browse offers the ability to transform raw Hi-C contact matrix into a (log) correlation matrix, whereas my5C can generate the expected Hi-C signal and the ratio of observed to expected Hi-C signal. Juicebox indirectly offers the ability to perform all these transformations through the Juicer software. Other tools require the user to externally apply the transformations to the raw Hi-C data prior to upload.

Several software tools are available to carry out these external transformations. Juicer is the complementary software package to Juicebox that can process sequencing reads from a Hi-C experiment into .hic files that contain contact matrices at multiple resolutions and in various transformations. HiC-Pro [40] offers similar capabilities to Juicer but uses a tab delimited sparse matrix to store the output, in both raw and matrix balanced forms. The HOMER suite of tools can generate dense Hi-C contact matrices and supports a rich set of downstream operations for transforming and analyzing Hi-C data [41]. Ay and Noble [9] provide a full review of Hi-C processing tools.

Lastly, we note that certain tools can visualize or compare multiple datasets simultaneously, a useful capability for investigating changes in 3D conformation of chromatin across different cell types or conditions. Juicebox and my5C can load two datasets, allowing the user to flip between heatmap visualizations and visualizing the ratio of Hi-C signals in the two data sets. The Epigenome Browser is able to visualize multiple Hi-C datasets, each as an individual track. The 3D Genome Browser and Hi-Browse currently support visualization of a single Hi-C dataset; however, we note that Hi-Browse offers a method to identify statistically significant differential regions based on edgeR [42].

## Complementary datasets

As we noted before, it is essential to plot genomic features together with Hi-C data when investigating, for example, the interplay between chromatin conformation and gene regulation. Because the Epigenome Browser and the 3D Genome Browser specialize in this task, these tools provide many publicly available datasets, primarily generated by the ENCODE and Roadmap Epigenomics consortia. Furthermore, many relevant annotation tracks of various genomic features (genes, GC islands, repeat regions) are available, offering a rich collection of features that can assist in interpretation of Hi-C data. Although Juicebox does not provide browser-like capabilities, the tool does offer a collection of genomic features, allowing a degree of joint visualization by placing tracks on the edges of the heat map visualization (Figure 1C). my5C tool generates links to the UCSC Genome Browser for loci of interest, allowing the user to seperately visualize other genomic features.

All the tools that offer visualization of genomic features—Juicebox, the Epigenome Browser, and the 3D Genome Browser—also support the capability to upload user genomic data, such as gene annotations or ChIP-seq peaks. Well defined standards for file formats for such data types are already in place. These formats include the BED file format that defines genomic features relative to genomic intervals, and wig and bedgraph formats that are used to store continuous signal along the length of the genome.

In addition to classic browser tracks, the 3D Genome Browser offers the ability to visualize two additional features that characterize 3D interactions: ChIA-PET and DNase-seq linkage annotations. ChIA-PET linkages are experimentally determined three dimensional contacts that are tethered by a specific protein [5], whereas DNase-seq linkages are predicted functional interactions between DNase hypersensitive sites [31]. These linkages are visualized as arcs and can aid in interpreting contacts revealed by a virtual 4C plot. For example, a virtual 4C plot focusing on the promoter of the NANOG gene displays a potential promoter-enhancer interaction upstream of the gene (Figure 2B).

## Implementation

In addition to differing in their functionality, the five tools that we examined differ fairly substantially in how they are implemented. In particular, although all of the tools are freely available, only HiBrowse, the Epigenome Browser, and Juicebox are open source. Furthermore, the Epigenome Browser and Juicebox can be installed to run on the user’s local computer, circumventing the need to access online servers through the internet. This can be desirable for analyses that require confidentiality or significant computational resources. Local installation for Juicebox only requires a 64-bit Java distribution, whereas installing the Epigenome Browser depends on multiple software packages and server services, described in detailed, step-by-step instructions in the corresponding manual.

All of the tools provide a graphical user interface that is available through a web browser interface or via Java Web Start, thus requiring no or minimal installation. Unless a local installation is performed, all tools also require an internet connection. Access to tools that use a web browser interface is available through any operating system. For local installations, the Epigenome browser supports Linux and MacOS operating systems.

All the tools come with documentation, although we note that the documentation of the 3D Genome Browser is curently being updated. The Epigenome Browser has its own wiki page that offers a set of how-to’s for creating and managing files for storing track information. Additionally, we note that Juicebox and the Epigenome browser have active online discussion groups that are maintained by the developers of these tools.

For each visualization tool, we profiled the speed of two important operations: loading user data and visualizing loci of sizes that are appropriate for both browser-based and heatmap-based tools (Table 2). Multiple factors, such as internet connection speed and server load, make it challenging to set up an exact benchmarking protocol; thus, we only report the approximate speed of loading operations, on the order of seconds, minutes or hours, and we report an average duration for visualization tasks. For benchmarking, we set the resolution parameter to either 40 kbp or 50 kbp, commonly used resolutions that strike a balance between sparsity and detail. We found that Juicebox, the Epigenome Browser and the 3D Genome Browser process user data in binary formats in a few seconds. HiBrowse and my5C do not support loading of a complete dataset at these resolutions, instead requiring the user to upload the HiC contact matrix corresponding to the region of interest. The average times required to visualize 1 Mb and 10 Mb heatmaps showed that tools that do not use a browser framework are faster, with Juicebox and my5C being the fastest tools. Browser-based tools are generally slower, especially for 10 Mb loci, consistent with the browser-based tools’ intended focus on local visualizations. We stress that user experience may differ from our benchmark due to differences in data sets, internet bandwidth and other parameters; thus, we offer this benchmark as a general guideline rather than an absolute measure of speed.

**Table 2:** Speed benchmarks for loading and visualizing Hi-C data.

| Tool | Loading User Data | Visualization of 1Mb Loci | Visualization of 10Mb Loci |
| --- | --- | --- | --- |
| Juicebox | seconds | 10 seconds | 13 seconds |
| Hi-Browse | NA | 10 seconds | 86 seconds |
| my5C | NA | 1 seconds | 3 seconds |
| 3D Genome Browser | seconds | 7 seconds | 43 seconds |
| Epigenome Browser | seconds | 33 seconds | 73 seconds |

## Discussion

Although all of the tools we have discussed aim to represent the same Hi-C data, some tools are better suited to understanding the conformation of chromatin at large or small scales. Hi-Browse and my5C are well equipped to visualize large scale conformations, like that of a complete genome or an individual chromosome. The Epigenome and 3D Genome browsers can better represent conformations at smaller scales, such as contacts involving a single gene, further enriching such visualization with other genomic features. Juicebox strikes a balance between these two approaches, offering browser-like functionality to visualize supplemental data next to a matrix-based Hi-C visualization. Thus, the tool of choice for a Hi-C analysis task depends on the nature of the inquiry regarding chromatin conformation. In this study, we offer two example uses cases to illustrate our point: browsers are very capable of probing effects of chromatin conformation on the regulation of a single gene (Figure 2), whereas heat maps are better suited to probing the overall organization of a single chromosome (Figure 1).

All the tools we review offer a graphical user interface and do not require programming skills to operate, making them broadly accessible. On the other hand, although it is easy to use these tools to create sophisticated visualizations of Hi-C data, processing and converting Hi-C data into the required contact matrix format requires at least a basic understanding of programming. None of the visualization tools we reviewed offer the ability to process raw Hi-C reads into a contact matrix, but other toolkits are available to automate such tasks (reviewed in [9]). In addition to the tools we reviewed here, software packages such as HiCplotter [43] and HiTC [44] offer visualization capabilities but require programming capabilities.

We have discussed visualization of raw or normalized Hi-C data, but other transformations of the data can be visualized using the same set of tools. For example, statistical confidence measures, such as p-values produced by methods like Fit-Hi-C [12] or diffHiC [45], can be converted to a contact matrix format and then visualized using the tools reviewed here. Hi-C data can also be used to infer the 3D structure of the chromatin (methods reviewed in [46]). The software tools reviewed here could be used to visualize the Euclidean distance matrix induced by such a 3D model. Direct visualization of the 3D models, especially in conjunction with other genomic features, is potentially very powerful. Several visualization tools for 3D genome structures are available, including GMol [47], Shrec3D [18], TADBit [48] and TADKit (http://sgt.cnag.cat/3dg/tadkit/demo/index.html#/project/browser).

## Acknowledgments

This work was funded by National Institutes of Health awards U54 DK107979 and U41 HG007000.

